# Anomalous brain energy in old age by wavelet analysis of ERP during a Stroop task

**DOI:** 10.1101/2021.01.11.426273

**Authors:** Sergio M. Sánchez-Moguel, Roman Baravalle, Sofía González-Salinas, Osvaldo A. Rosso, Thalía Fernández, Fernando Montani

**Author notes:** Both authors contributed equally to the current project (Sergio M. Sánchez-Moguel, Roman Baravalle). **Corresponding authors’ email addresses** (Fernando Montani), (Sergio M. Sánchez-Moguel).

## Abstract

By event-related potentials (ERP) during a counting Stroop task it was shown that the elderly with excess in theta activity in their electroencephalogram (EEG) are at risk of cognitive decline and have a higher neuronal activity during stimulus categorization than the elderly with a normal EEG. It was suggested that this increased neuronal activity could have a compensatory function. However, the quantification of energy associated with the enhanced neuronal activity was not investigated in this group. By wavelet analysis, we measured total and relative energy in ERP during the execution of a counting Stroop task in two groups of elderly: one with excess in theta activity (Theta-EEG, n = 23) and the other with normal EEG (Normal-EEG, n = 23). In delta, theta, and alpha bands, the Theta-EEG group used a higher amount of total energy as compared to the Normal-EEG group for both types of stimuli, interference and no interference. In theta and alpha bands, the total energy was higher in the Theta-EEG group, specifically in the window of 258-516 ms, coinciding with stimulus categorization. Given that no major behavioral differences were observed between EEG groups, we suggest that a higher energy in delta, theta, and alpha bands is one of the neurobiological mechanisms that allows the Theta-EEG group to cope with the cognitive demands of the task. However, this increased energy might not be an effective mechanism in the long term as it could promote a metabolic and cellular dysregulation that would trigger the transition to cognitive impairment.

**SIGNIFICANCE STATEMENT:** By using wavelet transform analysis we report that the elderly with excess in theta activity show a higher energy in delta, theta, and alpha bands during the categorization of stimuli in a counting Stroop task. Our findings imply that this increase neuronal activity might be related to a dysregulated energy metabolism in the elderly with theta excess that could explain the progress to cognitive impairment in this group. The analysis of energy by wavelet transform in data obtained by ERP complements other techniques that evaluate the risk of cognitive impairment.

## INTRODUCTION

Healthy aging is accompanied by a natural detriment of physical and cognitive abilities (Román Lapuente and Sánchez Navarro, 1998). In particular, inhibitory control (Thomas et al., 2010; Rey-Mermet and Gade, 2018) and attention (Thomas et al., 2010; Diamond, 2020) are importantly affected. Changes in brain electrical activity, which can be measured noninvasively by the EEG, are tightly related to the aforementioned cognitive processes (Buzsáki, 2006; Lopes da Silva, 2011). Some authors have proposed that changes in the EEG of the elderly, obtained under resting conditions, are not only the result of normal aging but can contain signs of undergoing subclinical pathologic processes (Chang et al., 2011). Moreover, excess in delta and theta band frequencies of resting EEG from healthy elderly, compared to a normative base according to age, is an excellent predictor of cognitive detriment in the following seven years (Prichep et al., 2006; van der Hiele et al., 2008). Recently we showed that healthy elderly with an excess of theta EEG activity are not only at risk of developing cognitive decline but already have impairments in inhibitory control processing at the electrophysiological level (Sánchez-Moguel et al., 2018).

Stroop tasks have been used during event-related potentials (ERP) and functional magnetic resonance imaging (fMRI) to study the decrease in the efficiency of inhibitory processing during healthy and pathological aging (West and Alain, 2000; Amieva, 2004; Kaufmann et al., 2008; Ramos-Goicoa et al., 2016; Sánchez-Moguel et al., 2018). An over-recruitment of neuronal activity during aging was observed using fMRI during the execution of Stroop tasks; this enhanced neuronal activity is proposed to have a compensatory function (Cabeza, 2002; Milham et al., 2002; Cabeza et al., 2004; Langenecker et al., 2004; Zysset et al., 2007; Mathis et al., 2009). Furthermore, fMRI studies showed higher brain activity in older people with mild cognitive impairment (MCI) compared to healthy elderly (Kaufmann et al., 2008).

In our earlier work, we proposed that the elderly with excess in theta EEG activity have an increased neuronal activity in ERP during a counting Stroop task (Sánchez-Moguel et al., 2018). We also suggested that the higher brain activity in ERP during a counting Stroop task was related to the categorization of stimuli, which would play a compensatory role. A higher neuronal activity is related to more energy; however, we have not quantified the energy associated with any 3 cognitive process in the elderly with excess in theta activity. As the elderly with excess in theta activity are probably in a previous stage of MCI, we hypothesize that they might already be having a dysregulation in brain energy, wich is a hallmark of neurodegenerative diseases (Mattson and Arumugam, 2018).

Wavelet transform (WT) can help us to know the amount of energy used during the execution of Stroop tasks. The main advantage of wavelet analysis over Fourier analysis is the optimal time-frequency resolution, then, we can follow the brain frequency dynamics over time (Rosso et al., 2006). The wavelet analysis allows us to have a standard frequency decomposition of EEG signals over time (Goupillaud et al., 1984; Rosso et al., 2005, 2006). This is a desirable property, because we can track the frequency changes of the EEG signal over time and detect at which time point of the Stroop task the maximum amount of energy occurs.

The general objective of this study was to explore, using WT, if there were differences in the amount of energy in ERP during the performance of a counting Stroop task between a group of elderly with an excess of theta activity in their EEG and a group of elderly with normal EEG. The specific objective was to evaluate the amount of energy between EEG groups for each of the frequency bands (i.e., delta, theta, alpha, beta, and gamma) across different time windows of the ERP. We expected to find higher energy in the group with theta excess, specifically in the time window associated with categorization of stimuli.

## MATERIALS AND METHODS

### Participants

Forty-six healthy older adults aged over 60 years were recruited to participate in the study (26 females). The inclusion criteria were to be right-handed, to have more than nine years of schooling, to have an average level of intelligence (Wechsler Intelligence Scale for adults 90-190, (Wechsler, 2003)), and to not have any psychiatric disorder according to their age (NEUROPSI, (Ostrosky-Solís et al., 1999)); Q-LES-Q questionnaire, > 70%, (Endicott et al., 1993); Mini-Mental State Examination, > 27, (Reisberg et al., 1982, 2008); Global Deterioration Scale, 1-2 (Reisberg et al., 1982, 2008); Alcohol Use Disorders Identification Test, < 5 (Babor et al., 2001); Beck Depression Inventory, < 4 (Beck et al., 1961); Geriatric Depression Scale, < 5 (Yesavage et al., 1982). Furthermore, subjects had no signs of chronic diseases such as diabetes or hypercholesterolemia. The subjects were classified into two groups according to the characteristics of their EEG. Subjects in the Normal-EEG group presented normal EEGs, from both the quantitative and qualitative points of view, and subjects in the Theta-EEG group presented an excess of theta activity for their age in at least one electrode; further described below. The project was approved by the bioethics committee of the Neurobiology Institute of the National Autonomous University of Mexico (UNAM). ERP analyses of the participants were published by Sánchez-Moguel et al. (2018) and are further analyzed here using wavelets.

### EEG analysis

Based on the next analysis, participants were classified as with a normal EEG (Normal-EEG group) or with excess in the theta band (Theta-EEG group); 23 subjects made up each group (13 females in each group).

The EEG from 19 tin electrodes (10-20 International System, ElectroCap™, International Inc.; Eaton, Ohio) referenced to linked ear lobes was recorded from each subject in the resting condition with eyes closed using a MEDICID ™*IV* system (Neuronic Mexicana, S.A.; Mexico) and Track Walker ™ v5.0 data system for 15 min. The EEG was digitized using the MEDICID IV System (Neuronic A.C.) with a sampling rate of 200 *Hz* using a band-pass filter of 0.5 – 50 *Hz*, and the impedance was kept below 5 *k*Ω. Twenty-four artifact-free segments of 2.56 *s* each were selected, and the quantitative EEG analysis was performed offline using the fast Fourier transform to obtain the power spectrum every 0.39 *Hz*; also the geometric power correction (Hernández et al., 1994) was applied, and absolute (AP) and relative power (RP) in each of the four classic frequency bands were obtained: Delta (1.5 - 3.5 *Hz*), theta (3.6 - 7.5 *Hz*), alpha (7.6 - 12.5 *Hz*), and beta (12.6 - 19 *Hz*). These frequency ranges were the same as those used for the normative database (Valdés et al., 1990) provided by MEDICID IV. Z-values were obtained for AP and RP, comparing subject’s values with values of the normative database [*Z* = (*x* - *μ*)/*σ*, where *μ* and *σ* are the mean value and the standard deviation of the normative sample of the same age as the subject, respectively]; *Z*-values > 1.96 were considered abnormal (*p* < 0.05).

### Counting Stroop task

In the counting Stroop task, subjects are asked to answer how many words are presented in a slide, regardless of the meaning of the word itself (Bush et al., 2006). Subjects increase their response times and tend to make more mistakes when the meaning of the word does not match the number of times that the word appears; this phenomenon is known as the Stroop or interference effect (MacLeod, 1991).

### Behavioral task

Series of one, two, three, or four words that denote numbers (“one,” “two,” “three,” “four”) were presented in the center of a 17-inch computer screen. Time presentation of the stimuli was 500 ms, and the interstimulus interval was 1,500 ms. An incongruent condition, herein referred as Interference stimulus, consisted of a trial where the number of presented words did not correspond with the meaning of the word. The congruent condition, further referred as No Interference stimulus, consisted of a trial in which the number of presented words and the meaning of the word that was presented matched. A total of 120 Interference and 120 No Interference stimuli were randomly presented.

Subjects were asked to indicate the number of times that the word appeared in each trial, using a response pad that they held in their hands. One-half of the participants used their left thumbs to answer “one” or “two” and their right thumbs to indicate “three” or “four”; the other half of the participants used their opposite hand to counterbalance the motor responses. The participants were asked to answer as quickly and accurately as possible. We ensured that the participants understood the instructions by presenting a brief practice task before the experimental session.

### ERP acquisition and analysis

The EEGs were recorded with 32 Ag/AgCl electrodes mounted on an elastic cap (Electrocap) while the participant performed the counting Stroop task, using NeuroScan SynAmps amplifiers (Compumedics NeuroScan) and the Scan 4.5 software (Compumedics NeuroScan). Electrodes were referenced to the right earlobe (A2), and the electrical signal was collected from the left earlobe (A1) as an independent channel. Recordings were re-referenced offline in two ways: (a) to the averaged earlobes, as was usually performed in previous studies, and (b) to the average reference. The EEG was digitized with a sampling rate of 500 Hz using a band pass filter of 0.01 to 100 Hz. Impedances were kept below 5 kΩ. An electrooculogram was recorded using a supraorbital electrode and an electrode placed on the outer canthus of the left eye.

ERP were obtained for each subject and experimental condition (i.e., No Interference and Interference). Epochs of 1,500 ms were obtained for each trial that consisted of 200-ms prestimulus and 1,300-ms post-stimulus intervals. An eye movement correction algorithm (Gratton et al., 1983) was applied to remove blinks and vertical ocular-movement artifacts. Low pass filtering for 50 Hz and a 6-dB slope was performed offline. A baseline correction was performed using the 200-ms pre-stimulus time window, and a linear detrend correction was performed on the whole epoch. Epochs with voltage changes exceeding ±80 μV were automatically rejected from the final average. The epochs were visually inspected, and those with artifacts were also rejected. Averaged waveforms for each subject and each stimulus type included only those trials that corresponded to correct responses.

### Wavelet transform and wavelet-based measures

The ERPs were next subjected to a wavelet analysis. Unlike Fourier analysis, in which the sine and cosine functions are used, the wavelet transform is based on functions that are vanishing oscillating functions (Rosso et al., 2006). Within the wavelet multiresolution decomposition framework, a wavelet family *ψ_a,b_* is a set of elemental functions generated by scaling and translating a unique admissible mother wavelet *ψ*(*t*):

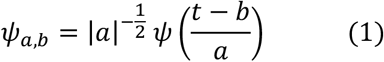

where 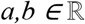, *a*≠0 are the scale and translation parameter, respectively, and *t* is the time (Rosso et al., 2006). In this paper we use the Daubechies 2 as a mother wavelet.

The continuous wavelet transform (CWT) of a signal 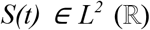 (the space of real square summable functions) is defined as the correlation between the signal *S*(*t*) with the family wavelet *ψ_a,b_* for each *a* and *b*:

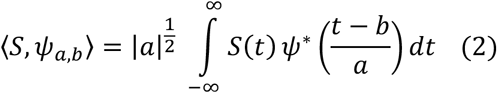

where * means complex conjugation. In principle, the CWT gives a highly redundant representation of the signal because it produces an infinite number of coefficients (Rosso et al., 2006). A nonredundant and efficient representation is given by the discrete wavelet transform (DWT), which also ensures complete signal reconstruction. For a special selection of the mother wavelet function *ψ*(*t*) and the discrete set of parameters *a_j_* = *2^-j^* and *b_j,k_* = *2^-j^ k*, with 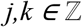, the family *ψ_j,k_*(*t*) = *2^j/2^ ψ*(*2^j^ t - k*) constitutes an orthonormal basis of 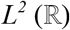. Any arbitrary function of this space can therefore be uniquely decomposed, and the decomposition can be inverted (Rosso et al., 2006). The wavelet coefficients of the DWT are 〈*S, ψ_j,k_*〉 = *C_j_*(*k*). The DWT produces only as many coefficients as there are samples within the signal under analysis *S*(*t*), without any loss of information.

Let us assume that the signal is given by the equally sampled values *S* = *{s_0_ (n), n = 1,…,M}*, with *M* being the total number of samples. If the decomposition is carried out over all resolution levels, *N_J_* = *log(M)*, the wavelet expansion reads:

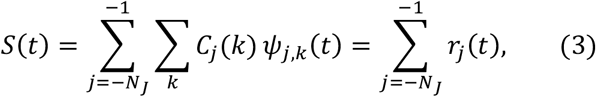

where the wavelet coefficients *C_j_*(*k*) can be interpreted as the local residual errors between successive signal approximations at scales *j* and *j-1*, respectively, and *r_j_*(*t*) is the detail signal at scale *j*, which contains information of the signal *S*(*t*) corresponding to the frequencies 2^j-1^ ω_s_ ≤ |ω| ≤ 2^j^ ω_s_, ω_s_ being the sampling frequency (Rosso et al., 2005, 2006).

Since the family *ψ_j,k_(t)* is an orthonormal basis for 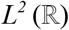, the concept of wavelet energy is similar to the Fourier theory energy. Thus, the energy at each resolution level, *j = −1,…,-N_J_*, will be the energy of the detail signal:

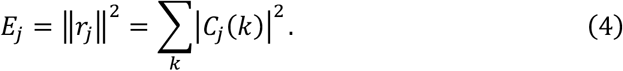

The total energy can be obtained summing over all the resolution levels

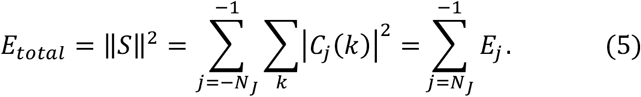

Finally, we define the relative wavelet energy (RWE) through the normalized *ρ_j_* values:

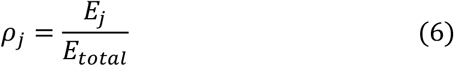

for the resolution levels *j = −1, −2,…,-N_J_*. The distribution *P^(W)^ ≡ {ρj}* can be viewed as a time-scale distribution, which is a suitable tool for detecting and characterizing phenomena in the time and frequency spaces (Rosso et al., 2006).

Another extension of this discrete wavelet transform is the discrete wavelet packet transform (DWPT). The DWPT is a generalization of the DWT that at level *j* of the transform partitions the frequency axis into *2^j^* equal width frequency bands, often labeled n = 0,…,2^j-1^. Increasing the transform level increases frequency resolution, but starting with a series of length *N*, at level *j* there are only *N / 2^j^* DWPT coefficients for each frequency band *n* (Percival and Walden, 2000).

The wavelet packets can be organized on an orthonormal basis of the space of finite energy signals. The main advantage of using wavelet packets is that the standard wavelet analysis can be extended with a flexible strategy. Thus the description of the given signal can be well adapted according to the significant structures (Blanco et al., 1998). The resulting DWPT yields what can be called a time-scale-frequency decomposition because each DWPT coefficient can be localized to a particular band of frequencies and a particular interval of time (Percival and Walden, 2000). Here we use the flexibility of the DWPT to combine the energy of the decomposition frequency bands, in order to have an insight into the typical clinical frequency band decomposition: Delta, theta, alpha, beta, and gamma. Finally, we have the energy {*E_Delta_, E_Theta_, E_Alpha_, E_Beta_, E_Gamma_*} corresponding to each band, and the relative energy {*ρ_Delta_, ρ_Theta_, ρ_Alpha_, ρ_Beta_, ρ_Gamma_*} for each one of the five bands. The energy corresponding to each band is obtained by adding all the values of *E_j_* for all the *j* that satisfy *2^j-1^ω_s_* ≤ |*ω*| ≤ *2^j^ω_s_, ω_s_* being the sampling frequency and |*ω*| being within the frequency interval corresponding to one of the five clinical frequency

### Statistical analysis

The behavioral data from the counting Stroop task, and the total and relative energy were analyzed using ANOVAs according to the variables of interest in each set of results. Repeated measures were included for Stimulus, Bands, and Windows, as required. A Tukey post hoc test was used to make comparisons among groups. Data were processed, analyzed, and plotted using R and Matlab.

## RESULTS

### Behavioral results of the counting Stroop task

**Table 2** shows the mean percentage of correct responses and response times (RT) by each group and type of stimulus. For the RT, there was no main effect of Group (F(1, 44) = 0.73, p = 0.3963), while Stimulus and the Group X Stimulus interaction were significant (Stimulus: F(1, 44) = 85.89, p < 0.0001; Group X Stimulus: F(1, 44) = 4.66, p = 0.0363). Post hoc analysis showed that RT for the Interference stimuli were larger than the response times for No Interference stimuli both within the Theta-EEG (Mean Difference (MD) = 42.61 ms, p < 0.001) and within the Normal-EEG groups (MD = 68.5 ms, p < 0.001); the Theta-EEG group showed fewer differences between stimulus types than the Normal-EEG group. There were no differences between groups for the same type of stimulus (Interference: p = 0.39, No Interference p = 0.99). We applied the arcsine to the percentage of correct responses in order to approximate the distribution of the data to a Gaussian distribution to use parametric statistical tests. We observed a significant main effect of Stimulus (F(1, 44) = 62.43, p < 0.0001) with a lower percentage of correct answers in the Interference than in the No Interference condition; however, there were no main effects of Group or Group X Stimulus interaction (Group: F(1, 44) = 0.09, p = 0.76, Group X Stimulus: F(1, 44) = 1.24, p = 0.27). These results showed that, at the behavioral level, the Theta-EEG and Normal-EEG groups showed a Stroop effect and that they answered similarly despite the differences in their resting EEG.

**Table 1.** Frequency for each band analyzed. The interval for each band is specified.

| Band | Frequency Interval (Hz) |
| --- | --- |
| Delta | [1.9531 - 3.9063) |
| Theta | [3.9063 - 7.8125) |
| Alpha | [7.8125 - 11.7188) |
| Beta | [11.7188 - 19.5313) |
| Gamma | [19.5313 - 39.0625) |

**Table 2.**
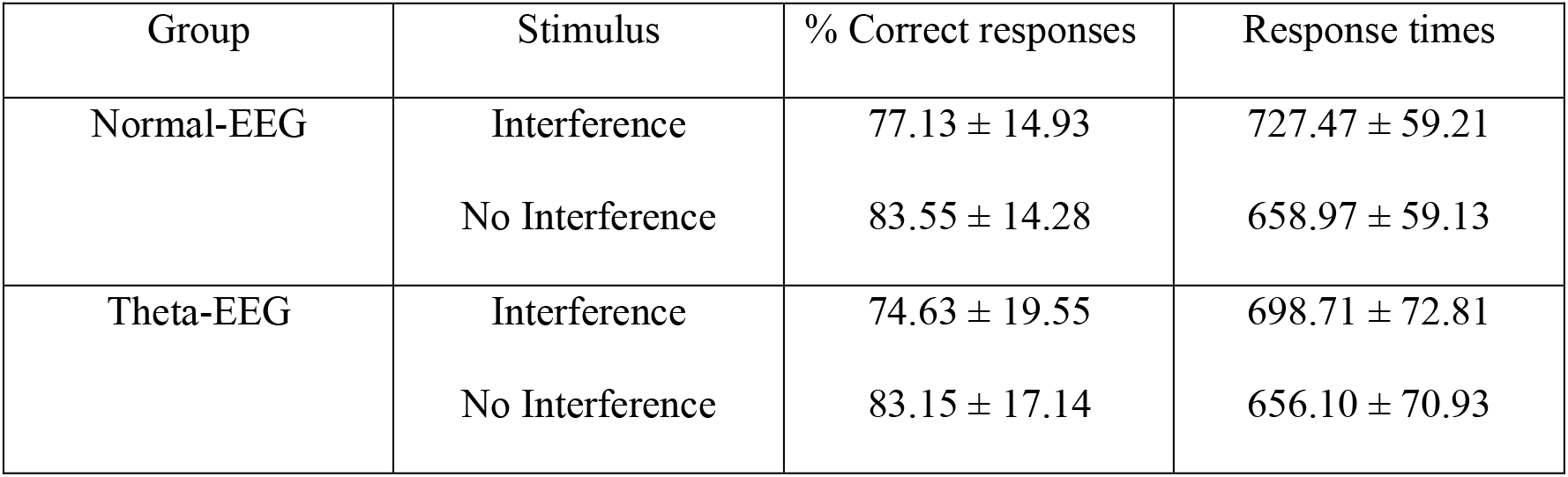
Behavioral performance during the counting Stroop task. Data are shown as Mean ± standard deviation (SD); response times are expressed in ms.

### Total energy

We first compared the total energy on each band between Theta-EEG and Normal-EEG groups, obtaining the total energy of the average of all electrodes (reference electrodes A1, A2 were discarded) and averaging across the counting Stroop trials for each type of stimulus. In **Figure 1**, the total energy for each group and stimulus type is shown in each of the frequency bands. For the delta band we found a main effect of Group (F(1, 86) = 11.003, p = 0.00133), while neither Stimulus (F(1, 86) = 0.416, p = 0.51961) nor the Group X Stimulus interaction were significant (F(1, 86) = 0.036, p = 0.84973). In the theta band, there was a main effect of Group (F(1, 86) = 11.605, p = 0.001), while no significant differences were observed in Stimulus (F(1, 86) = 0.031, p = 0.862) or in the Group X Stimulus interaction (F(1, 86) = 0.01, p = 0.919). Similarly, in the alpha band there was a main effect of Group (F(1, 86) = 8.539, p = 0.00444), while Stimulus (F(1, 86) = 0.002, p = 0.96375) and the Group X Stimulus interaction remained without statistical significance (F(1, 86) = 0.004, p = 0.95266).

**Figure 1.**
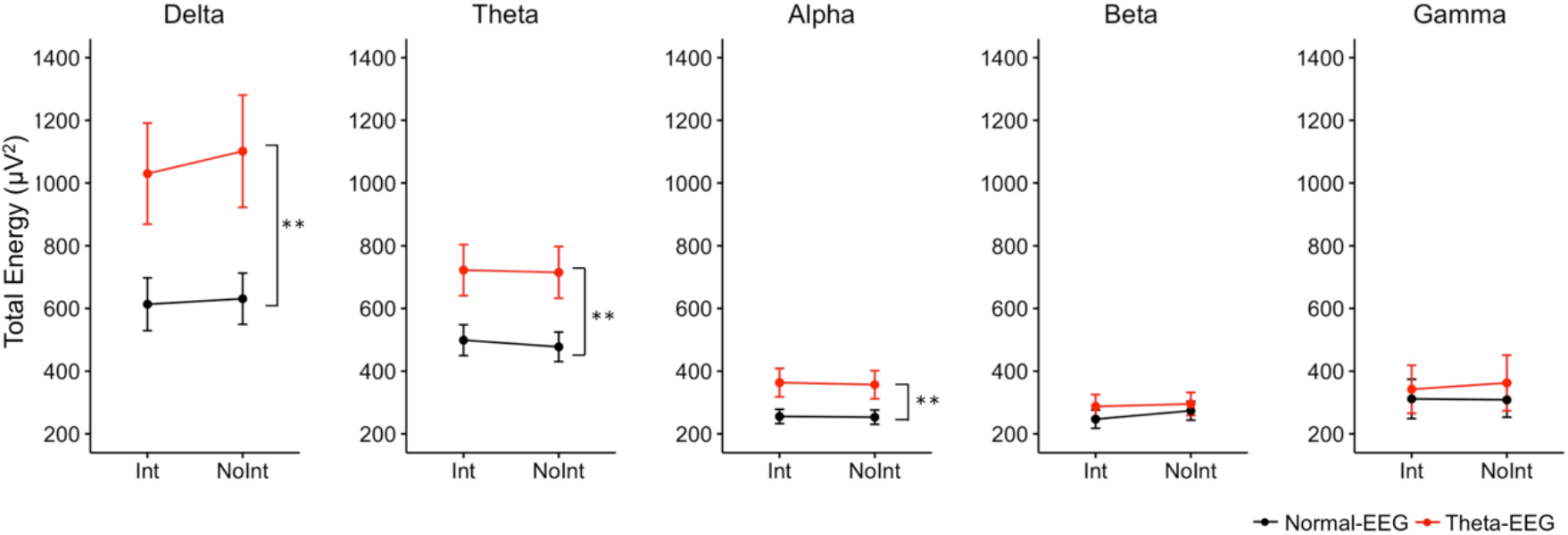
Total energy. Data were obtained from the average across electrodes (reference electrodes excluded) for Interference (Int) and No Interference (NoInt) stimuli during the counting Stroop task. ** p < 0.01 for group factor from two-way ANOVA. Data are expressed as means with standard error bars.

For the beta band, neither Group (F(1, 86) = 0.836, p = 0.363) nor Stimulus (F(1, 86)=0.099, p=0.753) or the Group X Stimulus interaction were significant (F(1,86) = 0.078, p = 0.781). Similar results were observed in the gamma band, no significant differences were found for Group (F(1, 86) = 0.330, p = 0.567), Stimulus (F(1, 86) = 0.127, p = 0.723) or the Group X Stimulus interaction (F(1, 86) = 0.027, p = 0.869).

Altogether, our analysis of the total energy showed a higher amount of energy in the Theta-EEG group in the delta, theta, and alpha bands irrespective of the type of stimulus presented during the counting Stroop task. In contrast, no significant differences in the total energy were observable in the beta and gamma bands, **Figure 1**.

We explored how the total energy was distributed among the electrodes for each band, **Figure 2**. We observed a higher total energy in the Theta-EEG group than in the Normal-EEG group in delta and theta bands. This increase in total energy was similar for both types of stimulus. For the delta band, the total energy increase was located in the midline electrodes, while for the theta band, the total energy increase was more pronounced in occipital electrodes. No increase in total energy was visible in alpha, beta, and gamma bands.

**Figure 2.**
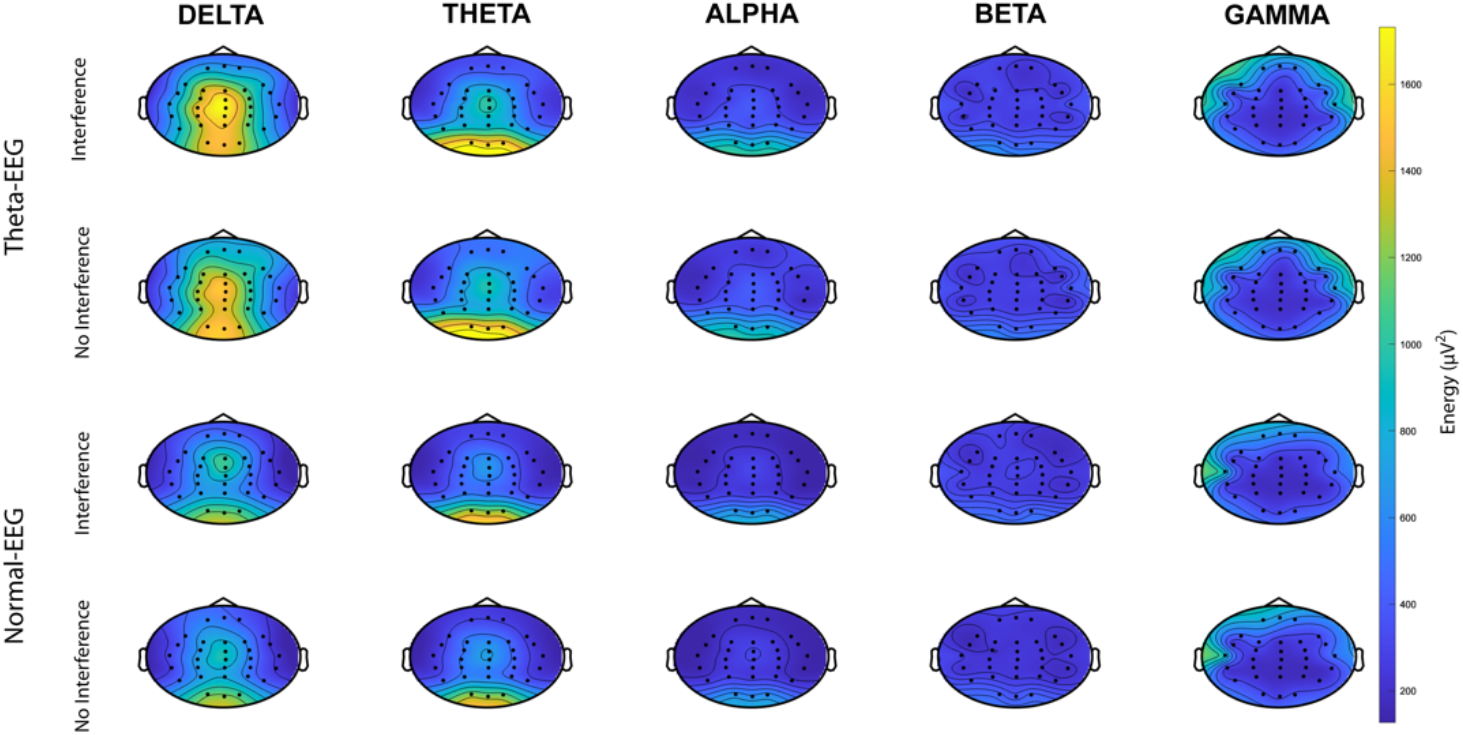
Topographic distribution of total energy. Total energy is shown for the different bands during Interference and No Interference stimuli according to the EEG group. The color scale is expressed in μV^2^.

### Relative energy

To consider variations in the total amount of energy among subjects, we further studied the relative wavelet energy for the entire signal. The relative energy corresponds to the amount of energy in a given band, relative to the total energy involved in all bands.

In **Figure 3** the relative energy per frequency band is shown for each group and stimulus type. In the delta band we observed a main effect of Group (F(1, 86) = 10.346, p = 0.00183) but not of Stimulus (F(1, 86) = 0.678, p = 0.41269) or the Group X Stimulus interaction (F(1, 86)=0.056, p = 0.81351). In the theta band neither of the variables nor the interaction between them were significant [Group (F(1, 86) =1.651, p = 0.202); Stimulus (F(1, 86)=0.165, p = 0.686); Group X Stimulus (F(1, 86) = 0.186, p = 0.668)]. For the alpha band there were no statistical differences for any of the effects or the interaction between them [Group (F(1, 86) = 0.070, p = 0.793); Stimulus (F(1, 86) = 0.021, p = 0.886); Group X Stimulus (F(1, 86) = 0.062, p = 0.804)].

**Figure 3.**
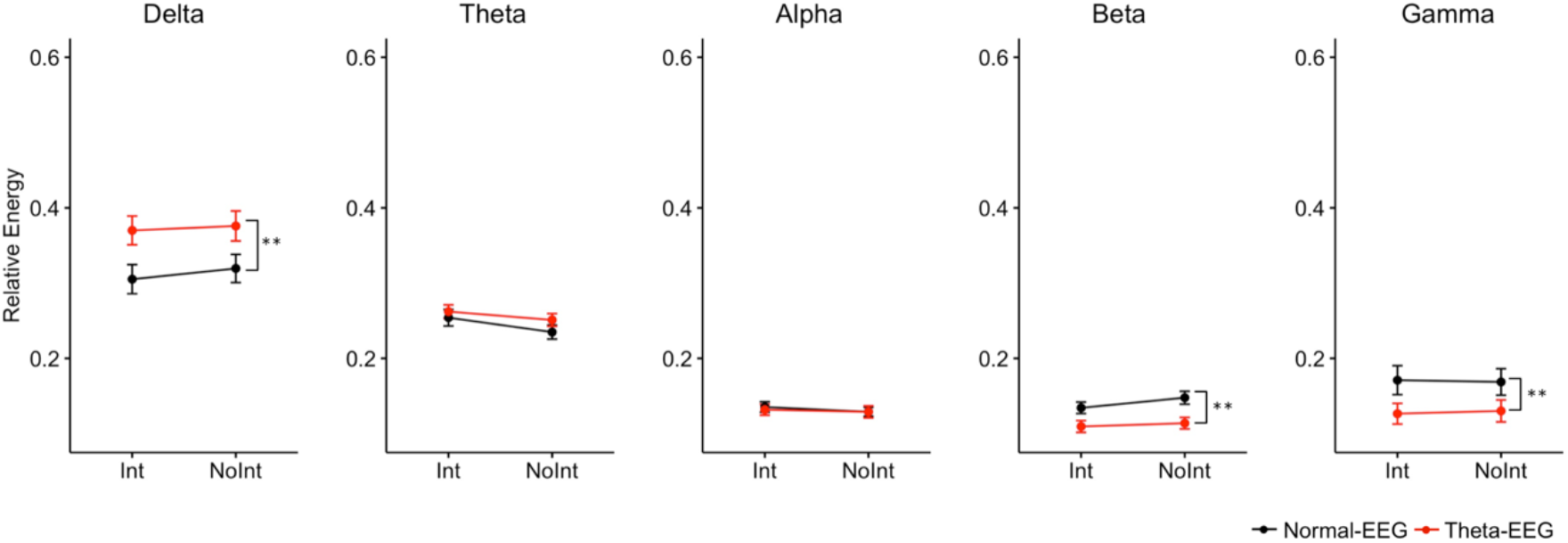
Relative energy. Data are shown for each band according to the EEG group and to the type of stimulus. Data are expressed as means with standard error bars. ** p < 0.01 for group factor from two way-ANOVA.

The relative energy in the beta band showed a main effect of Group (F(1, 86) = 13.401, p = 0.000433) but not for Stimulus (F(1, 86) = 1.676, p = 0.198983) or for the Group X Stimulus interaction (F(1, 86) = 0.312, p = 0.577787). Similarly, for the gamma band, there was a main effect of Group (F(1, 86)=7.017, p = 0.00961), but no differences were observed for Stimulus (F(1, 86) = 0.259, p = 0.61188) or for the Group X Stimulus interaction (F(1, 86) = 0.038, p = 0.84581).

Our analysis thus showed that even after normalizing by the total amount of energy used during the task, the Theta-EEG group showed an increase of energy in the delta band, as compared to the Normal-EEG group, which was independent of the type of stimulus presented. Interestingly, a decrease in relative energy was observed in the beta and gamma bands.

### Total energy across windows

To analyze the signal in the time-space, we took time windows of at least 2^7^ + 1 = 129 points, which corresponded to 258 ms; this procedure allowed us to analyze five time-windows in the ERP signal. **Figure 4** shows the total energy per window. For the delta band, significant main effects of Group (F(1, 430) = 21.058, p = 5.86e-06) and Window (F(4, 430) = 12.610, p = 1.04e-09) were observed but there were not significant effects in Stimulus (F(1, 430) = 0.149, p = 0.7) or the interaction among factors [Group X Stimulus (F(1, 430) = 0.062, p = 0.804); Group X Window (F(4, 430) = 0.79, p = 0.532); Stimulus X Window (F(4, 430) = 0.307, p = 0.873); Group X Stimulus X Window (F(4, 430) = 0.367, p = 0.832)]. The analysis of the Window factor showed that the total energy in the window 258-516 ms was higher than the total energy from windows 0-258, 516-774, and 774-1032 ms (p ≤ 0.03783 for each comparison). For this same band, the total energy in the window 1032-1290 was higher than the energy in the window 0-258 (p = 0.00893).

**Figure 4.**
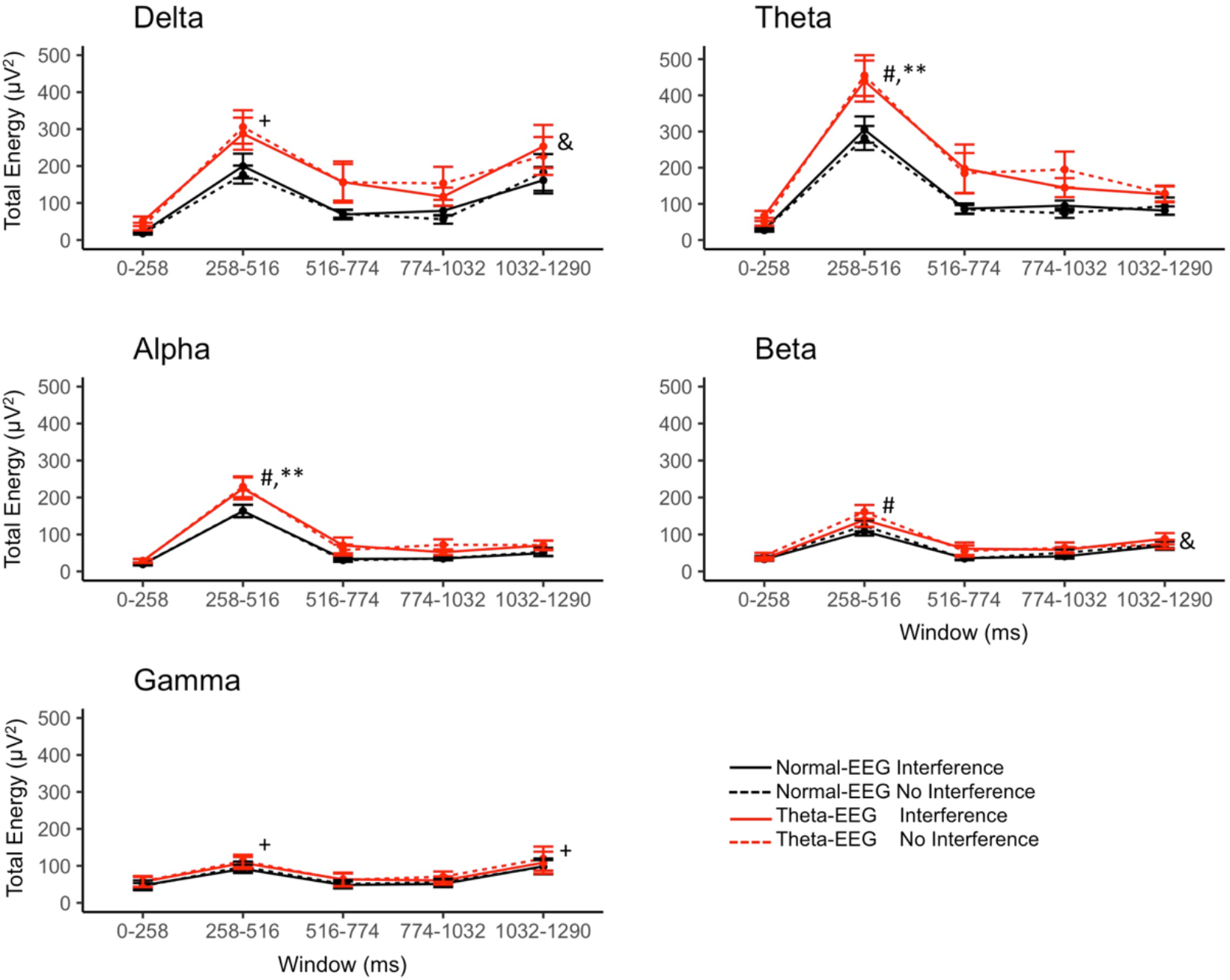
Total energy for time windows and bands. Post hoc test of Group X Window: p < 0.01 between Normal-EEG and Theta-EEG for the 258-516 window. Post hoc test of Window: ^#^p < 0.05 compared to the windows 0-258, 516-774, 774-1032, and 1032-1290; ^+^p < 0.05 compared to 0-258, 516-774, and 774-1032 windows; ^&^p < 0.05 compared to 0-258 window. Data are expressed as means with standard error bars.

For the theta band, there were significant main effects of Group (F(1, 430) = 30.596, p = 5.53e-08) and Window (F(4, 430) = 17.546, p = 2.38e-13) but not of Stimulus (F(1, 430) = 0.279, p = 0.5978). The interaction Group X Window was close to significance (F(4, 430) = 2.229, p = 0.0651), while the other interactions were not significant [Group X Stimulus (F(1, 430) = 0.241, p = 0.624); Stimulus X Window (F(4, 430) = 0.298, p = 0.8794); Group X Stimulus X Window (F(4, 430) = 0.321, p = 0.8639)]. The analysis of the Window factor showed that the total energy in the window 258-516 ms was higher than the total energy in all the other windows (p < 0.0001 for all the comparisons). The post hoc test for the interaction Group X Window showed that in the window 258-516 ms the total energy was higher in the Theta-EEG group as compared to the Normal-EEG group (p = 0.0065), **Figure 4**.

For the alpha band, the total energy showed significant main effects of Group (F(1, 430) = 23.548, p = 1.71e-06) and Window (F(4, 430) = 34.484, p < 2e-16), while Stimulus was not significant (F(1, 430) = 0.005, p = 0.9452). The interaction Group X Window was close to statistical significance (F(4, 430) = 2.163, p = 0.0724), while other interactions did not show significant differences [Group X Stimulus (F(1, 430) = 0.03, p = 0.8622); Stimulus X Window (F(4, 430) = 0.053, p = 0.9947); Group X Stimulus X Window (F(4, 430) = 0.149, p = 0.9633)]. The analysis of the Window factor showed that the total energy in the window 258-516 ms was higher than the energy in all the other windows (p < 0.0001 for each comparison). The analysis of the interaction Group X Window showed that the total energy in the window 258-516 ms was higher in the Theta-EEG group as compared to the energy in the Normal-EEG group (p = 0.0079), **Figure 4**.

For the beta band we found significant main effects of Group (F(1, 430) = 10.868, p = 0.00106) and Window (F(4, 430) = 17.402, p = 3.03e-13), while there were no statistical differences for Stimulus (F(1, 430) = 0.311, p = 0.57723). None of the interactions of the factors was significant either [Group X Stimulus (F(1, 430) = 0.105, p = 0.74663); Group X Window (F(4, 430) = 1.046, p = 0.3829); Stimulus X Window (F(4, 430) = 0.295, p = 0.88137); Group X Stimulus X Window (F(4, 430) = 0.2, p = 0.93804)]. The post hoc comparisons of the Window factor showed that the total energy in 258-516 ms was higher than the energy in all the other windows (p ≤ 0.0143 for all the comparisons). Additionally, the energy in the 1032-1290 ms window was higher than the energy observed in the 0-258 ms window (p = 0.0261), **Figure 4**.

Finally, the analysis of total energy in the gamma band revealed a significant main effect of Window (F(4, 430) = 4.018, p = 0.00328), while Group (F(1, 430) = 3.242, p = 0.07246) and Stimulus (F(1, 430) = 0.318, p = 0.57321) did not reach significance. None of the interactions among factors was significant [Group X Stimulus (F(1, 430) = 0.053, p = 0.81849); Group X Window (F(4, 430) = 0.021, p = 0.99915); Stimulus X Window (F(4, 430) = 0.093, p = 0.98453); Group X Stimulus X Window (F(4, 430) = 0.037, p = 0.99741)]. The analysis of the Window factor showed that the total energy in the 258-516 ms window was higher than the energy in the windows 0-258, 516-774, and 774-1032 ms (p < 0.01 for all comparisons). The total energy in the window 1032-1290 ms was also higher than the energy observed in the windows 0-258, 516-774, and 774-1032 ms (p ≤ 0.002 for all comparisons), **Figure 4**.

Taken together, our analysis across windows revealed a higher amount of total energy in the Theta-EEG group as compared to the Normal-EEG group in the delta, theta, alpha, and beta bands irrespective of the type of stimulus presented. For theta and alpha bands, the total energy was higher in the Theta-EEG group than in the Normal-EEG group, specifically for the window 258-516 ms, **Figure 4**.

A topographical analysis of the total energy across windows for the theta band corroborated the relevance of the 258-516 ms window. The topographical distribution of the energy per group and type of stimulus through the time is shown in **Figure 5**. We observed a higher total energy in the Theta-EEG group as compared to the Normal-EEG group only in the 258-516 ms window. The amount of total energy looked similar for both types of stimulus in the same EEG group. The increased energy for this theta band in the Theta-EEG group was more prominent in mid-line and occipital electrodes. Similar changes were observed in delta and alpha bands (**Figure 5-1** and **5-2**): increased total energy for both types of stimulus in the Theta-EEG group was observed in mid-line and occipital electrodes towards frontal regions; this change was prominent in the 258-516 ms window.

**Figure 5.**
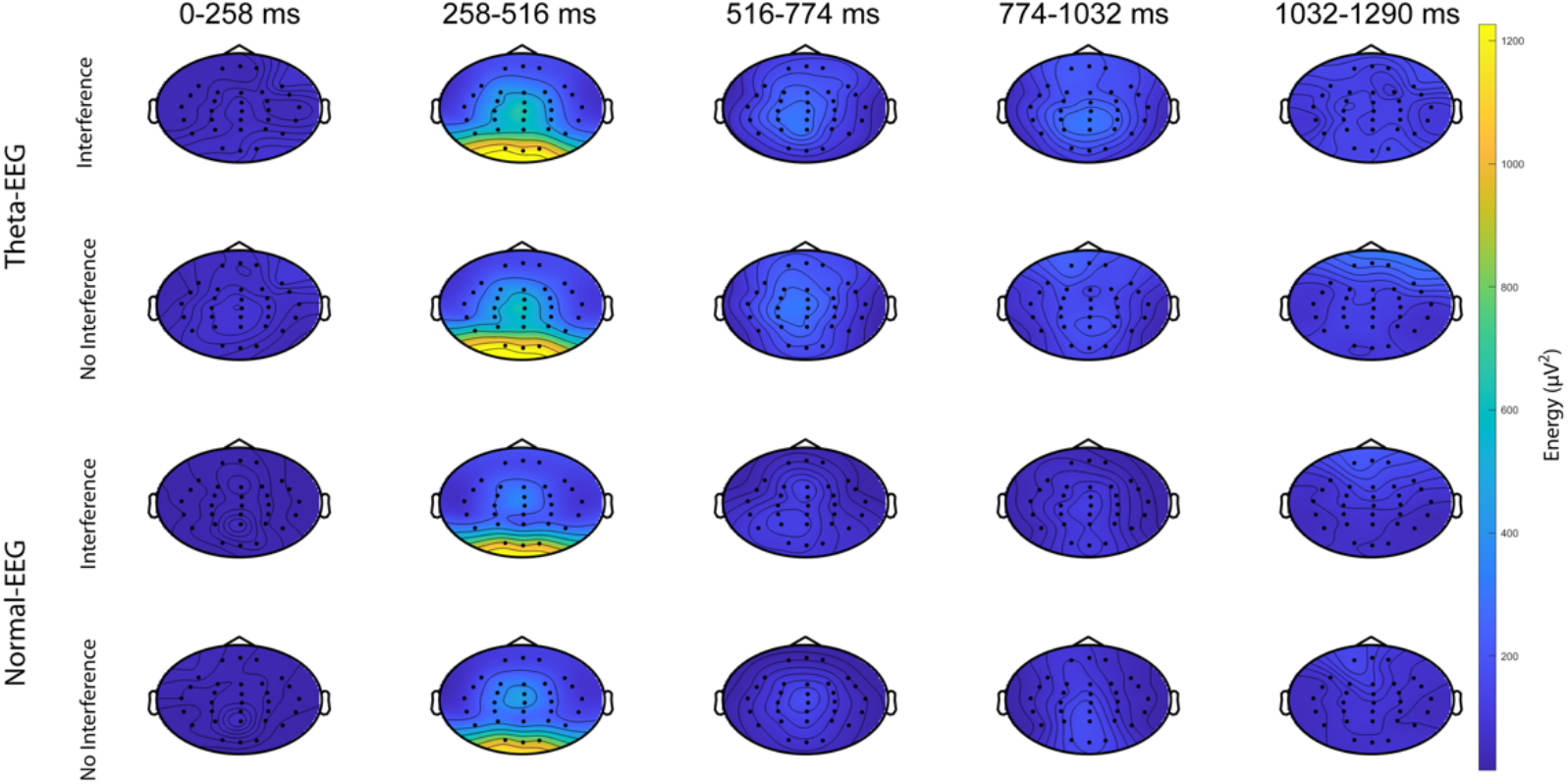
Topographic distribution of the total energy in the theta band. Data are shown for each window and according to the group and to the type of stimulus. The color scale is expressed in μV^2^.

**Figure 5-1.**
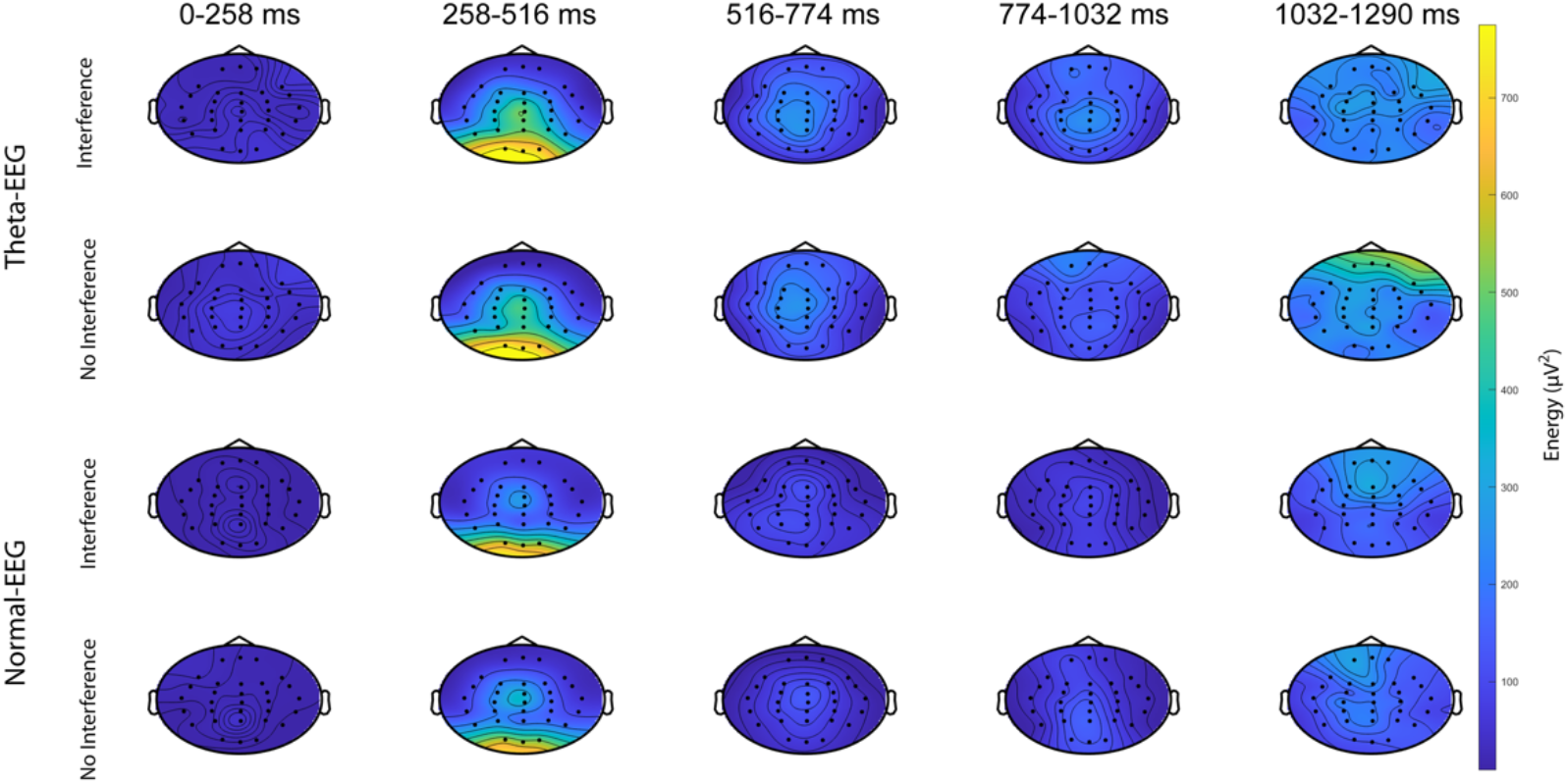
Topographic distribution of the total energy in the delta band. Data are shown for each window and according to the group and to the type of stimulus. The color scale is expressed in μV^2^.

**Figure 5-2.**
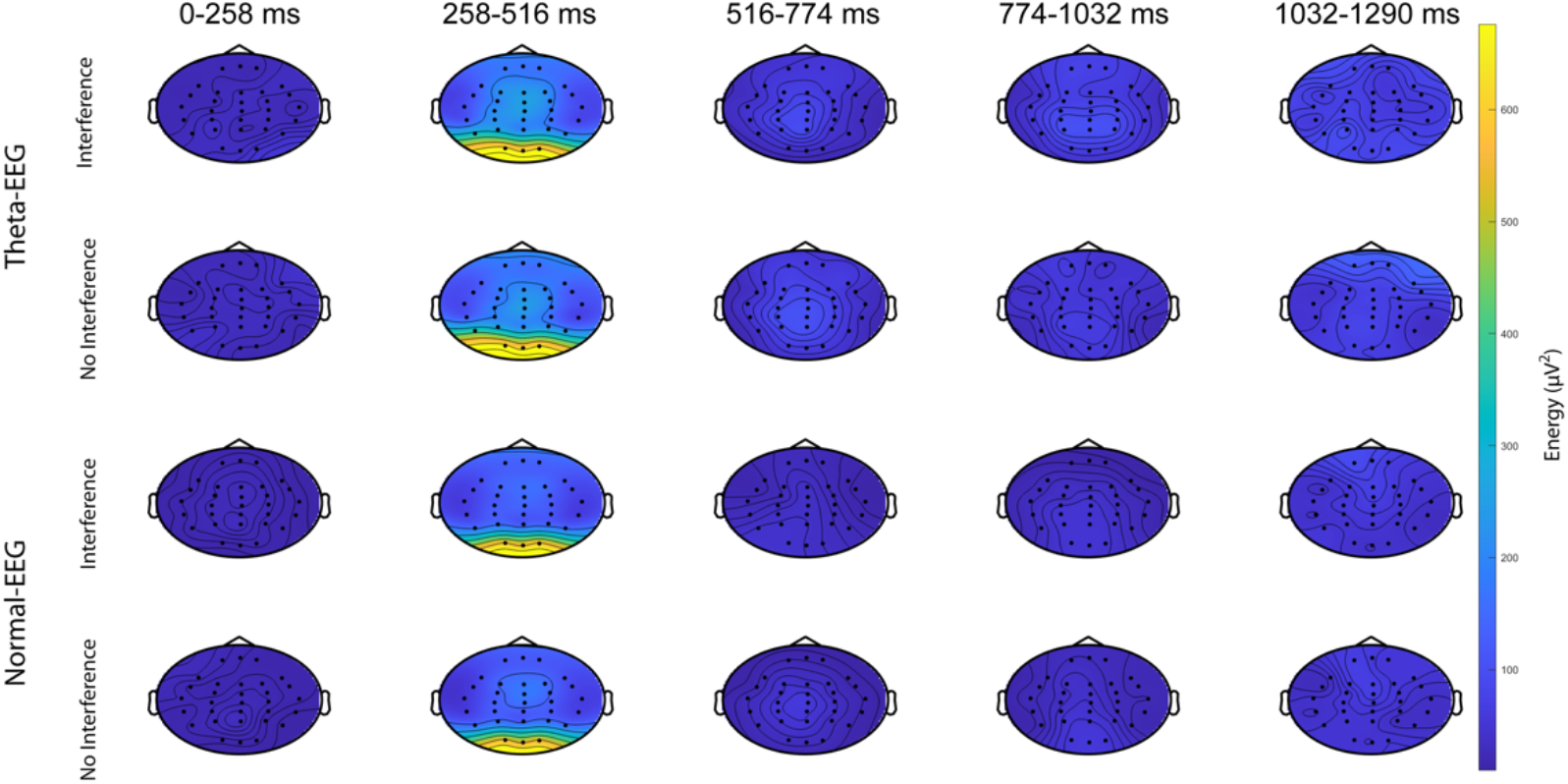
Topographic distribution of the total energy in the alpha band. Data are shown for each window and according to the group and to the type of stimulus. The color scale is expressed in μV^2^.

### Wavelet analysis on central electrodes

Then, we performed an analysis for central electrodes to better depict the information added by wavelet analysis when studying the data obtained by ERP, **Figure 6**. The wavelet transform of the voltage in the CPZ electrode showed a significant Group X Window interaction in delta, theta, and alpha bands in the total energy, **Table 3**. The post hoc test showed that the total energy was higher in the Theta-EEG group than in the Normal-EEG group in the 258-516 ms window for the three bands (p ≤ 0.0141). The differences in total energy for CPZ electrode in delta, theta, and alpha bands were independent of the type of stimulus presented (Stimulus, Group X Stimulus, Stimulus X Window, and Group X Stimulus X Window were not significant; **Table 3**). Although the Group X Window interaction was significant for the beta band (**Table 3**), the post hoc comparison did not show any statistical difference between Theta-EEG and Normal-EEG conditions for any given window (p ≥ 0.0720). For the gamma band, there was a significant effect of Group and Window but not for other factors or interactions, **Table 3**.

**Figure 6.**
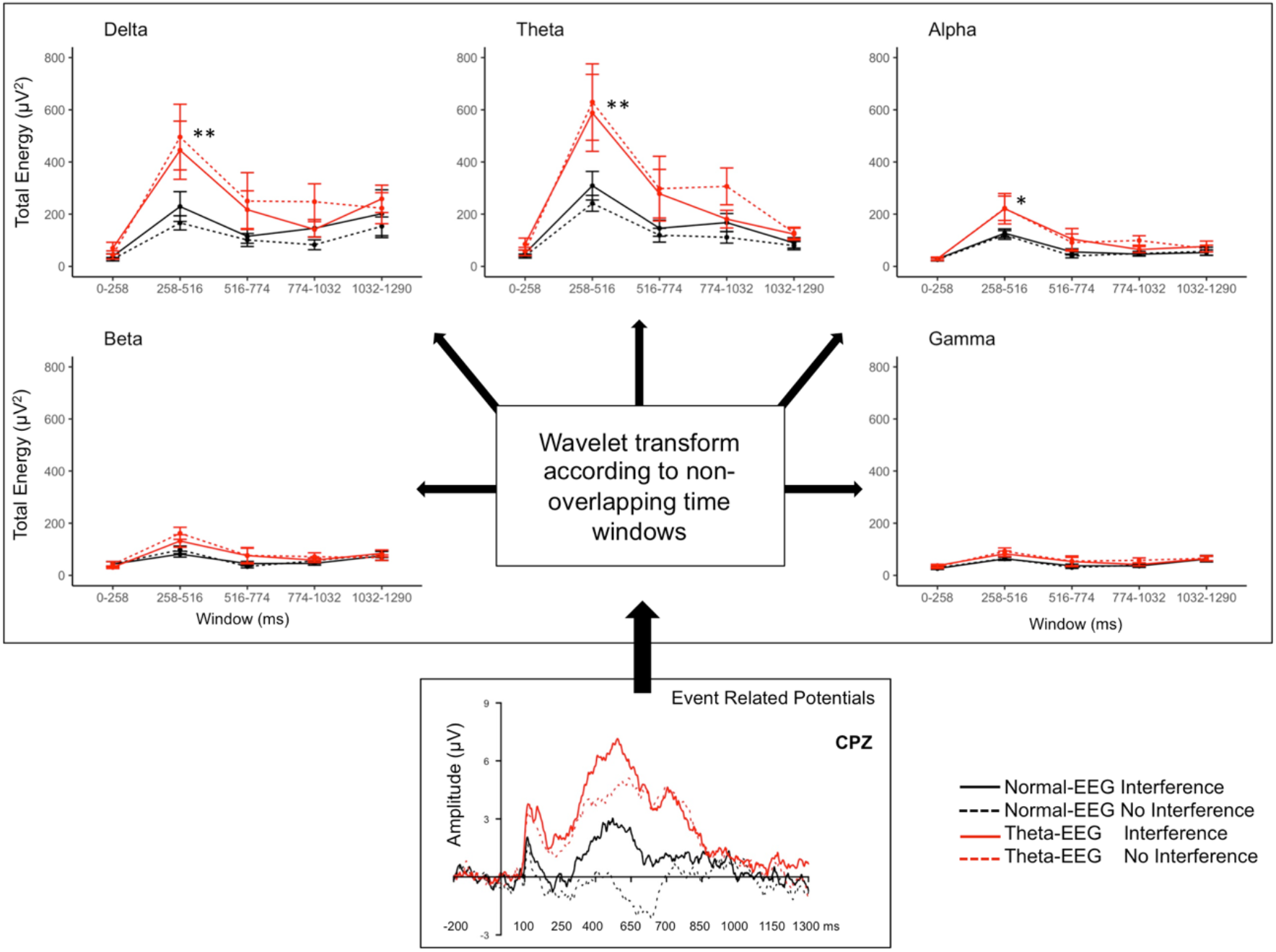
Total energy in the CPZ electrode. The amplitude of the signal obtained by event-related potentials during a counting Stroop task was further analyzed by wavelet transform. **p<0.01, *p <0.05 for the window 258-516 ms when comparing Theta-EEG versus Normal-EEG in the post hoc analysis of the interaction Group X Window. No significant differences were observed for other windows or for the type of stimulus.

**Table 3.** Two-way ANOVA results for the total energy in central electrodes. There were main effects of Group and Window for all bands in all electrodes. The Group X Window interaction was significant for alpha in more anterior electrodes while for delta and theta bands this effect was observed in more posterior electrodes. Post hoc results for the Group X Window interaction are described in the text.

| Electrode | Band/ ANOVA factors | Delta | Theta | Alpha | Beta | Gamma |
| --- | --- | --- | --- | --- | --- | --- |
|  |  | F(1,430);<br>p-value | F(1,430);<br>p-value | F(1,430);<br>p-value | F(1,430);<br>p-value | F(1,430);<br>p-value |
| FCZ | Group | 9.1; 0.003 | 12.5; 0.0004 | 12.5; 0.0004 | 7.1; 0.008 | 4.2; 0.04 |
|  | Stimulus | 0.003; 0.9 | 0.3; 0.6 | 0.006; 0.9 | 1.5; 0.2 | 0.8; 0.4 |
|  | Window | 5.9; 0.0001 | 9.7; 1.54e-07 | 19.2; 1.64e-14 | 13.0; 5.21e-10 | 7.4; 8.72e-06 |
|  | Group X Stimulus | 0.2; 0.7 | 0.6; 0.4 | 0.3; 0.6 | 0.3; 0.6 | 0.09; 0.8 |
|  | Group X Window | 0.5; 0.8 | 1.3; 0.3 | 2.4; 0.05 | 2.0; 0.1 | 0.4; 0.8 |
|  | Stimulus X Window | 1.0; 0.4 | 0.5; 0.7 | 0.2; 0.97 | 0.8; 0.5 | 0.9; 0.4 |
|  | Group X Stimulus X Window | 0.4; 0.8 | 0.4; 0.8 | 0.3; 0.9 | 0.4; 0.8 | 0.4; 0.8 |
| CZ | Group | 11.7; 0.0007 | 14.6; 0.0002 | 12.8; 0.0004 | 6.5; 0.01 | 6.2; 0.01 |
|  | Stimulus | 0.1; 0.8 | 0.2; 0.6 | 0.01; 0.9 | 1.3; 0.3 | 0.7; 0.4 |
|  | Window | 5.6; 0.0002 | 9.5; 2.42e-07 | 16.9; 6.73e-13 | 12.9; 6.49e-10 | 8.7; 9.07e-07 |
|  | Group X Stimulus | 1.1; 0.3 | 1.1; 0.3 | 0.2; 0.7 | 0.01; 0.9 | 0.1; 0.7 |
|  | Group X Window | 0.9; 0.5 | 1.8; 0.1 | 2.6; 0.03 | 2.4; 0.05 | 0.5; 0.8 |
|  | Stimulus X Window | 0.6; 0.6 | 0.6; 0.7 | 0.1; 0.97 | 0.4; 0.8 | 1.1; 0.4 |
|  | Group X Stimulus X Window | 0.4; 0.8 | 0.4; 0.8 | 0.2; 0.99 | 0.3; 0.9 | 0.3; 0.9 |
| CPZ | Group | 16.9; 4.62e-05 | 20.8; 6.75e-06 | 15.2; 0.0001 | 8.2; 0.005 | 9.3; 0.002 |
|  | Stimulus | 0.3; 0.61 | 0.3; 0.6 | 0.02; 0.9 | 0.8; 0.4 | 0.3; 0.6 |
|  | Window | 4.5; 0.001 | 8.5; 1.40e-06 | 16.2; 2.8e-12 | 11.9; 3.45e-09 | 8.8; 8.3e-07 |
|  | Group X Stimulus | 1.4; 0.2 | 1.3; 0.3 | 0.2; 0.8 | 0.04; 0.8 | 0.2; 0.6 |
|  | Group X Window | 2.5; 0.0447 | 3.7; 0.006 | 2.7; 0.03 | 3.1; 0.02 | 0.9; 0.5 |
|  | Stimulus X Window | 0.5; 0.8 | 0.4; 0.8 | 0.1; 0.98 | 0.2; 0.95 | 0.96; 0.4 |
|  | Group X Stimulus<br>X Window | 0.4; 0.8 | 0.4; 0.8 | 0.1; 0.97 | 0.3; 0.91 | 0.2; 0.95 |
| PZ | Group | 19.9; 1.02e-05 | 24.6; 1.04e-06 | 14.4; 0.0002 | 5.9; 0.02 | 8.2; 0.005 |
|  | Stimulus | 0.4; 0.6 | 0.2; 0.6 | 0.003; 0.96 | 0.6; 0.4 | 0.3; 0.6 |
|  | Window | 5.2; 0.00042 | 8.3; 1.96e-06 | 13.9; 1.12e-10 | 9.8; 1.46e-07 | 7.2; 1.24e-05 |
|  | Group X Stimulus | 0.7; 0.4 | 0.99; 0.3 | 0.03; 0.9 | 0.04; 0.8 | 0.2; 0.7 |
|  | Group X Window | 3.8; 0.0047 | 4.9; 0.0007 | 1.9; 0.1 | 2.2; 0.07 | 0.8; 0.5 |
|  | Stimulus X<br>Window | 0.2; 0.9 | 0.3; 0.9 | 0.1; 0.9 | 0.1; 0.9 | 0.7; 0.6 |
|  | Group X Stimulus<br>X Window | 0.5; 0.7 | 0.4; 0.8 | 0.2; 0.9 | 0.2; 0.9 | 0.3; 0.9 |

The total energy in the PZ electrode showed a significant Group X Window interaction for the delta, and theta bands, **Table 3**. The total energy was higher in the Theta-EEG group for the 258-516 ms window (p ≤ 0.0005 for delta and theta bands). In the alpha, beta, and gamma bands, only the factors Group and Window showed statistical significance, **Table 3**.

A similar increase in the total energy in the alpha band was observed in FCZ and CZ electrodes for the Theta-EEG group. There was a significant Group X Window interaction in the alpha band for both electrodes, **Table 3**. The total energy was higher in the Theta-EEG group than in the Normal-EEG group only in the 258-516 ms window (FCZ p = 0.0288; CZ p = 0.0175). For the CZ electrode, the Group X Window interaction was significant for the beta band; however, the post hoc comparisons did not show statistical differences for any specific time window, **Table 3**. For FCZ and CZ electrodes, only the factors Group and Window showed statistical significance for delta, theta, and gamma bands, **Table 3**.

Taken together, our analysis of wavelet transform for central electrodes showed that the total energy was higher in the Theta-EEG group than in the Normal-EEG group in the 258-516 ms window in the delta and theta bands for more posterior electrodes (CPZ, PZ). The increase in total energy in the Theta-EEG group was observed in the 258-516 ms window for the alpha band in more anterior electrodes (FCZ, CZ).

## DISCUSSION

In this study we aimed to explore whether the amount of energy obtained from ERP during a counting Stroop task was different between a group of elderly with an excess of theta activity in their EEG and a group of elderly with normal EEG. We evaluated the amount of energy between EEG groups for each of the frequency bands across different time windows. Overall, we found a higher energy in the group with theta excess that might help understand the increased risk of cognitive decline in this group of elderly.

### Behavioral evidence

The results at the behavioral level showed that there were no major differences between the groups. These were expected results if we consider that the only difference between the groups was at the electrophysiological level in the quantitative EEG analysis. Furthermore, the Theta-EEG and Normal-EEG groups showed a Stroop effect (i.e., longer reaction times for interference stimuli) and they answered with similar efficacy despite the differences in their resting EEG, **Table 2**.

### Wavelet evidence

#### Total energy analysis

During the counting Stroop task performance, we observed that, for both types of stimulus, the Theta-EEG group requires a higher energy in delta, theta, and alpha bands than the Normal-EEG group, **Figures 1 and 2**. Given that the total energy (μV^2^) is related to the number of synchronized active neurons and that no major differences between groups were observed in the performance of the counting Stroop task, we think that a higher expenditure of energy is the biological mechanism that allows the Theta-EEG group to cope with the cognitive demands of this task.

In fMRI it has been observed that the healthy elderly have a greater neural activity during the performance of Stroop tasks as compared to young subjects (Cabeza, 2002; Milham et al., 2002; Langenecker et al., 2004; Zysset et al., 2007; Mathis et al., 2009). This enhanced activity might be interpreted as a compensatory mechanism developed by them to achieve an optimum performance (Zysset et al., 2007; Mathis et al., 2009) or it may reflect a difficulty in recruiting specialized neuronal circuits (Cabeza, 2002). Furthermore, in fMRI and ERP studies, the elderly with MCI or with electrophysiological risk for cognitive decline, exhibited greater brain activity than healthy elderly (Kaufmann et al., 2008; Sánchez-Moguel et al., 2018). From the point of view of the performance of the task, it is evident that this compensatory mechanism is being effective in both the affected elderly with MCI and the Theta-EEG group. However, a higher energy expenditure of unspecialized neuronal circuits in the task might trigger anomalous cellular processes that are hallmarks of neurodegenerative diseases (Mattson and Arumugam, 2018), making this compensatory mechanism ineffective in the long term. The affected elderly will then have an anomalous activation of the involved circuits, and they will show a dysregulated energetic metabolism (Mattson and Arumugam, 2018).

The greater expenditure of energy in the Theta-EEG group observed in delta and theta bands agrees with the finding that increased activity in these bands predicts the development of cognitive impairment (Prichep et al., 2006; van der Hiele et al., 2008), **Figures 1, 2, and 4**. On the other hand, some studies suggest that increases in alpha power are related to success in inhibiting irrelevant information (Herrmann and Knight, 2001; Werkle-Bergner et al., 2012). This set of works supports our interpretation that the Theta-EEG group has a higher expenditure of alpha energy in order to perform the task with the same efficiency as the Normal-EEG group. Furthermore, the greater expenditure of energy in the alpha band in the Theta-EEG group can be explained by a topographic reorganization of the alpha rhythm during aging in which it is biased towards more frontal regions (Evans & Abarbanel, 1999), **Figure 5-2**. As mentioned earlier, these EEG changes are exacerbated in patients with dementia or MCI (Prichep et al., 1994).

We know that the brain resists entropy or disorder by maintaining its balance through the process of homeostasis (Friston et al., 2006; Friston, 2009, 2010). It is clear that the Theta-EEG group allocates more energy to maintain this homeostatic balance. However, over time the increased energy expenditure can promote neural metabolic imbalances more rapidly, causing the development of cognitive impairment.

#### Analysis of relative energy

There was a greater energy expenditure in both EEG groups in the delta and theta bands compared to the other bands, **Figure 3**. In the Theta-EEG group, the energy expenditure of the delta band was greater than in the Normal-EEG group; this relationship reverts in beta and gamma bands, **Figure 3**. Patients at risk of cognitive impairment (Prichep et al., 2006; van der Hiele et al., 2008) or that transition from MCI to Alzheimer (Huang et al., 2000; Jelic et al., 2000; Rossini et al., 2006) show an increase in the delta and theta power and a decrease in the beta relative power. The beta band is sensitive to the discrimination of interference and no interference stimuli in Stroop tasks (Schack et al., 1999), while the gamma band has a prominent role in the coupling of excitatory and inhibitory neuronal networks (Fries, 2009). This lower energy expenditure in beta and gamma bands in addition to the increased theta activity in the Theta-EEG group could then explain the inhibitory control impaired at the electrophysiological level previously reported by Sánchez-Moguel et al. (2018). Based on the higher risk of cognitive impairment of the Theta-EEG group (Sánchez-Moguel et al., 2018), we suggest that the greater relative energy in delta band and the lower relative energy in beta and gamma bands during the performance of the Stroop task may be related to the progression to MCI.

#### Analysis of total energy across time windows

The greater total energy in both EEG groups occurred in the 258-516 ms window for all bands, **Figures 4, 5, and 6**. In ERP studies, it has been observed that this time window is sensitive to the categorization of interference and no interference words (Zurrón et al., 2009; Sánchez-Moguel et al., 2018). Then we interpret that this greater energy expenditure is required to categorize the stimuli. The total energy for this time window was higher for the Theta-EEG group in the theta and alpha bands. We interpret that this increased energy is a mechanism that allows the Theta-EEG group to discriminate the stimuli with a similar efficiency as the Normal-EEG group.

The total energy expenditure for each stimulus condition in the different bands was similar within each EEG group, **Figures 1, 6, and Table 3**. This is an interesting result given the increased complexity of the interference as compared to the no interference stimuli because reading and counting processes are in competition (West and Alain, 2000; Bush et al., 2006) causing the RT to be longer in interference stimuli. We expected that longer RT would be related to higher energy expenditure. These results then suggest that in elderly adults, the processing of Interference and No Interference stimuli demands similar neuronal resources that might differ from young adults, a proposal that needs further study.

## OVERVIEW

In summary, the expenditure of energy was higher in the Theta-EEG group during a counting Stroop task. The energy analysis of ERP using wavelets showed that during the execution of the Stroop task: (1) Theta-EEG group assigns a greater amount of total energy in delta, theta, and alpha bands than the Normal-EEG group. (2) Theta-EEG group demands a higher amount of relative energy in delta band but less energy in beta and gamma bands than the normal-EEG group. (3) Theta-EEG group uses higher total energy in all-time windows in the delta, theta, alpha, and beta bands. (4) In the theta and alpha bands, the energy is greater in the Theta-EEG group, specifically in the time window 258-516 ms related to stimulus categorization processing. Thus the current findings emphasize the relevance of a wavelet analysis for diagnosis of neurological disorders, as in recent studies (Faust et al., 2015; Bhattacharyya and Pachori, 2017; Alturki et al., 2020).

We propose that this excessive energy expenditure in the Theta-EEG group is due because more neurons are recruited in order to perform the task with the same efficiency as the Normal-EEG group. However, we do not know if this energy expenditure is an effective long-term mechanism since neurons could be being recruited from unspecialized regions, and there could be cellular and metabolic imbalances that promote progress to cognitive impairment. Furthermore, since the Theta-EEG group participants have a higher risk of developing cognitive impairment and already show detriment of inhibitory control at the electrophysiological level, we suggest that this excessive energy expenditure begins to be anomalous.

Imaging techniques such as fMRI, diffusion tensor imaging, and magnetic resonance spectroscopy, that evaluate the neural networks involved in the task and metabolic expenditure, would complement our findings. Additionally, we suggest exploring energy expenditure during the performance of tasks related to other cognitive processes that are known to be altered in patients at risk of cognitive impairment.

## Acknowledgments

The authors acknowledge Leonor Casanova, Lourdes Lara, Graciela Alatorre-Cruz, and Teresa Alvarez for administrative support; Juan Silva-Pereyra, Erick Pasaye, Ramón Martínez, and Héctor Belmont for technical assistance; Marbella Espino, MD, for performing the neurological and psychiatric assessment. The first draft of the present work started during the summer school LACONEU, 2017; Sergio M. Sánchez-Moguel and Roman Baravalle received Fellowships to attend this event.

## REFERENCES

Alturki FA, AlSharabi K, Abdurraqeeb AM, Aljalal M (2020) EEG Signal Analysis for Diagnosing Neurological Disorders Using Discrete Wavelet Transform and Intelligent Techniques. Sensors 20:2505.

Amieva H (2004) Evidencing inhibitory deficits in Alzheimer’s disease through interference effects and shifting disabilities in the Stroop test. Arch Clin Neuropsychol 19:791–803.

Babor TF, Higgins-Biddle JC, Saunders JB, Monteiro MG (2001) AUDIT: the Alcohol Use Disorders Identification Test?: guidelines for use in primary health care. World Health Organ Available at: https://apps.who.int/iris/handle/10665/67205.

Beck AT, Ward CH, Mendelson M, Mock J, Erbaugh J (1961) An inventory for measuring depression. Arch Gen Psychiatry 4:561–571.

Bhattacharyya A, Pachori RB (2017) A Multivariate Approach for Patient-Specific EEG Seizure Detection Using Empirical Wavelet Transform. IEEE Trans Biomed Eng 64:2003–2015.

Blanco S, Figliola A, Quiroga RQ, Rosso OA, Serrano E (1998) Time-frequency analysis of electroencephalogram series. III. Wavelet packets and information cost function. Phys Rev E 57:932–940.

Bush G, Whalen PJ, Shin LM, Rauch SL (2006) The counting Stroop: a cognitive interference task. Nat Protoc 1:230–233.

Buzsáki G (2006) Rhythms of the brain. Oxford?; New York: Oxford University Press.

Cabeza R (2002) Hemispheric asymmetry reduction in older adults: the HAROLD model. Psychol Aging 17:85–100.

Cabeza R, Daselaar SM, Dolcos F, Prince SE, Budde M, Nyberg L (2004) Task-independent and task-specific age effects on brain activity during working memory, visual attention and episodic retrieval. Cereb Cortex N Y N 1991 14:364–375.

Chang BS, Schomer DL, Niedermeyer E (2011) “Normal EEG and sleep: adults and elderly. In: Niedermeyer’s Electroencephalography: Basic Principles, Clinical Applications, and Related Fields, pp 183–214. New York: Wolters Kluwer, Lippincott Williams & Wilkins.

Diamond A (2020) Chapter 19 - Executive functions. In: Handbook of Clinical Neurology (Gallagher A, Bulteau C, Cohen D, Michaud JL, eds), pp 225–240. Elsevier. Available at: http://www.sciencedirect.com/science/article/pii/B9780444641502000204.

Endicott J, Nee J, Harrison W, Blumenthal R (1993) Quality of Life Enjoyment and Satisfaction Questionnaire: a new measure. Psychopharmacol Bull 29:321–326.

Evans JR, Abarbanel A (1999) Introduction to Quantitative EEG and Neurofeedback. Elsevier.

Faust O, Acharya UR, Adeli H, Adeli A (2015) Wavelet-based EEG processing for computer-aided seizure detection and epilepsy diagnosis. Seizure 26:56–64.

Fries P (2009) Neuronal gamma-band synchronization as a fundamental process in cortical computation. Annu Rev Neurosci 32:209–224.

Friston K (2009) The free-energy principle: a rough guide to the brain? Trends Cogn Sci 13:293–301.

Friston K (2010) The free-energy principle: a unified brain theory? Nat Rev Neurosci 11:127–138.

Friston K, Kilner J, Harrison L (2006) A free energy principle for the brain. J Physiol-Paris 100:70–87.

Goupillaud P, Grossmann A, Morlet J (1984) Cycle-octave and related transforms in seismic signal analysis. Geoexploration 23:85–102.

Gratton G, Coles MGH, Donchin E (1983) A new method for off-line removal of ocular artifact. Electroencephalogr Clin Neurophysiol 55:468–484.

Hernández JL, Valdés P, Biscay R, Virues T, Szava S, Bosch J, Riquenes A, Clark I (1994) A global scale factor in brain topography. Int J Neurosci 76:267–278.

Herrmann CS, Knight RT (2001) Mechanisms of human attention: event-related potentials and oscillations. Neurosci Biobehav Rev 25:465–476.

Huang C, Wahlund L, Dierks T, Julin P, Winblad B, Jelic V (2000) Discrimination of Alzheimer’s disease and mild cognitive impairment by equivalent EEG sources: a cross-sectional and longitudinal study. Clin Neurophysiol Off J Int Fed Clin Neurophysiol 111:1961–1967.

Jelic V, Johansson SE, Almkvist O, Shigeta M, Julin P, Nordberg A, Winblad B, Wahlund LO (2000) Quantitative electroencephalography in mild cognitive impairment: longitudinal changes and possible prediction of Alzheimer’s disease. Neurobiol Aging 21:533–540.

Kaufmann L, Ischebeck A, Weiss E, Koppelstaetter F, Siedentopf C, Vogel SE, Gotwald T, Marksteiner J, Wood G (2008) An fMRI study of the numerical Stroop task in individuals with and without minimal cognitive impairment. Cortex 44:1248–1255.

Langenecker SA, Nielson KA, Rao SM (2004) fMRI of healthy older adults during Stroop interference. NeuroImage 21:192–200.

Lopes da Silva FH (2011) Neurocognitive processes and the EEG/MEG. In: Niedermeyer’s electroencephalography: Basic principles, clinical applications, and related fields, eds. Schomer, D. L., and Lopes da Silva, F.H., 6th ed., pp 1083–1112. New York, NY: Wolters Kluwer, Lippincott Williams & Wilkins.

MacLeod CM (1991) Half a century of research on the Stroop effect: an integrative review. Psychol Bull 109:163–203.

Mathis A, Schunck T, Erb G, Namer IJ, Luthringer R (2009) The effect of aging on the inhibitory function in middle-aged subjects: a functional MRI study coupled with a color-matched Stroop task. Int J Geriatr Psychiatry 24:1062–1071.

Mattson MP, Arumugam TV (2018) Hallmarks of Brain Aging: Adaptive and Pathological Modification by Metabolic States. Cell Metab 27:1176–1199.

Milham MP, Erickson KI, Banich MT, Kramer AF, Webb A, Wszalek T, Cohen NJ (2002) Attentional control in the aging brain: insights from an fMRI study of the Stroop task. Brain Cogn 49:277–296.

Ostrosky-Solís F, Ardila A, Rosselli M (1999) NEUROPSI: A brief neuropsychological test battery in Spanish with norms by age and educational level. J Int Neuropsychol Soc 5:413–433.

Percival DB, Walden AT (2000) Wavelet Methods for Time Series Analysis. Cambridge University Press.

Prichep LS, John ER, Ferris SH, Rausch L, Fang Z, Cancro R, Torossian C, Reisberg B (2006) Prediction of longitudinal cognitive decline in normal elderly with subjective complaints using electrophysiological imaging. Neurobiol Aging 27:471–481.

Prichep LS, John ER, Ferris SH, Reisberg B, Almas M, Alper K, Cancro R (1994) Quantitative eeg correlates of cognitive deterioration in the elderly. Neurobiol Aging 15:85–90.

Ramos-Goicoa M, Galdo-Álvarez S, Díaz F, Zurrón M (2016) Effect of Normal Aging and of Mild Cognitive Impairment on Event-Related Potentials to a Stroop Color-Word Task. J Alzheimers Dis 52:1487–1501.

Reisberg B, Ferris SH, de Leon MJ, Crook T (1982) The Global Deterioration Scale for assessment of primary degenerative dementia. Am J Psychiatry 139:1136–1139.

Reisberg B, Ferris SH, Kluger A, Franssen E, Wegiel J, de Leon MJ (2008) Mild cognitive impairment (MCI): a historical perspective. Int Psychogeriatr 20:18–31.

Rey-Mermet A, Gade M (2018) Inhibition in aging: What is preserved? What declines? A metaanalysis. Psychon Bull Rev 25:1695–1716.

Román Lapuente F, Sánchez Navarro JP (1998) Cambios neuropsicológicos asociados al envejecimiento normal. An Psicol 14:27–43.

Rossini PM, Del Percio C, Pasqualetti P, Cassetta E, Binetti G, Dal Forno G, Ferreri F, Frisoni G, Chiovenda P, Miniussi C, Parisi L, Tombini M, Vecchio F, Babiloni C (2006) Conversion from mild cognitive impairment to Alzheimer’s disease is predicted by sources and coherence of brain electroencephalography rhythms. Neuroscience 143:793–803.

Rosso OA, Martin MT, Figliola A, Keller K, Plastino A (2006) EEG analysis using wavelet-based information tools. J Neurosci Methods 153:163–182.

Rosso OA, Martin MT, Plastino A (2005) Evidence of self-organization in brain electrical activity using wavelet-based informational tools. Phys Stat Mech Its Appl 347:444–464.

Sánchez-Moguel SM, Alatorre-Cruz GC, Silva-Pereyra J, González-Salinas S, Sanchez-Lopez J, Otero-Ojeda GA, Fernández T (2018) Two Different Populations within the Healthy Elderly: Lack of Conflict Detection in Those at Risk of Cognitive Decline. Front Hum Neurosci 11 Available at: https://www.ncbi.nlm.nih.gov/pmc/articles/PMC5768990/.

Schack B, Chen AC, Mescha S, Witte H (1999) Instantaneous EEG coherence analysis during the Stroop task. Clin Neurophysiol Off J Int Fed Clin Neurophysiol 110:1410–1426.

Thomas AK, Dave JB, Bonura BM (2010) Theoretical Perspectives on Cognitive Aging. In: Handbook of Medical Neuropsychology: Applications of Cognitive Neuroscience (Armstrong CL, Morrow L, eds), pp 297–313. New York, NY: Springer New York. Available at: https://doi.org/10.1007/978-1-4419-1364-7_16.

Valdés P, Biscay R, Galán L, Bosch J, Zsava S, Virués T (1990) High resolution spectral EEG norms topography. Brain Topogr 3:281–282.

van der Hiele K, Bollen EL, Vein AA, Reijntjes RH, Westendorp RG, van Buchem MA, Middelkoop HA, van Dijk JG (2008) EEG markers of future cognitive performance in the elderly. J Clin Neurophysiol 25:83–89.

Wechsler D (2003) Wais-III escala wechsler de inteligencia para adultos-III, 2nd ed. México: Manual Moderno.

Weisz J, Czigler I (2006) Age and novelty: Event-related brain potentials and autonomic activity. Psychophysiology 43:261–271.

Werkle-Bergner M, Freunberger R, Sander MC, Lindenberger U, Klimesch W (2012) Inter-individual performance differences in younger and older adults differentially relate to amplitude modulations and phase stability of oscillations controlling working memory contents. NeuroImage 60:71–82.

West R, Alain C (2000) Age-related decline in inhibitory control contributes to the increased stroop effect observed in older adults. Psychophysiology 37:179–189.

Yesavage JA, Brink TL, Rose TL, Lum O, Huang V, Adey M, Leirer VO (1982) Development and validation of a geriatric depression screening scale: a preliminary report. J Psychiatr Res 17:37–49.

Zurrón M, Pouso M, Lindín M, Galdo S, Díaz F (2009) Event-related potentials with the stroop colour-word task: timing of semantic conflict. Int J Psychophysiol 72:246–252.

Zysset S, Schroeter ML, Neumann J, Yves von Cramon D (2007) Stroop interference, hemodynamic response and aging: An event-related fMRI study. Neurobiol Aging 28:937–946.

